# Cooperation between a root fungal endophyte and host-derived coumarin scopoletin mediates *Arabidopsis* iron nutrition

**DOI:** 10.1101/2025.06.04.657782

**Authors:** Lara Van Dijck, Dario Esposto, Charlotte Hülsmann, Milena Malisic, Anthony Piro, Ricardo F. H. Giehl, Gerd U. Balcke, Alain Tissier, Jane E. Parker

## Abstract

- Iron acquisition is a critical challenge for plants, especially in iron-deficient soils. Recent research underscores the importance of root-exuded coumarins in modulating the root microbiome community structure and facilitating iron uptake. However, interactions between root-associated fungi and coumarins in plant iron nutrition remain unknown. We investigated the mechanism by which a fungal endophyte, *Macrophomina phaseolina* (F80), enhances *Arabidopsis* iron nutrition.
- Fungal–coumarin interactions were assessed by profiling metabolites and measuring iron mobilisation in F80 cultures supplemented with specific coumarins, alongside quantifying growth performance and iron content in *Arabidopsis* coumarin-biosynthesis mutants inoculated with F80.
- Our findings reveal that an interaction between the coumarin scopoletin and F80 in the rhizosphere rescues plant growth under iron-limiting conditions by resolving the iron mobility bottleneck. F80 exhibits a capacity to modify scopoletin into iron-chelating catechol coumarin esculetin, thereby releasing available iron.
- We conclude that *Arabidopsis*-produced scopoletin functions as a precursor for fungal conversion into iron-chelating coumarins. By extending the role of coumarins from bacterial to fungal members of the root microbiota, this study places coumarins at the centre of commensal-mediated enhancement of plant iron nutrition across microbial kingdoms.

## Introduction

Plants have evolved sophisticated mechanisms to shape their root microbiomes. Microbial community structure in the rhizosphere is influenced by various root-exuded compounds, including a diverse class of secondary metabolites known as coumarins (Stassen *et al*., 2021). Coumarins are derived from the phenylpropanoid pathway and are produced by a wide range of dicotyledonous plants (Rajniak *et al*., 2018). Their broad biological activities which include anti-microbial, anti-oxidant, and anti-inflammatory properties, have attracted attention in plant and medical sciences (Borges *et al*., 2005). Coumarins can be found in above- and below- ground plant tissues (Stringlis *et al*., 2019; Robe *et al*., 2021a). Over 50 years ago, foliar coumarin accumulation was linked to pathogen defence (Zaynab *et al*., 2024), while more recent studies have highlighted their central role in plant iron acquisition at the root surface (Schmid *et al*., 2014; Sisó-Terraza *et al*., 2016; Rosenkranz *et al*., 2021).

Iron is an essential micronutrient for almost all life forms including plants and their associated microbes (Aznar *et al*., 2015). However, despite its high abundance in soil, iron is often bio-unavailable to soil-inhabiting organisms due to poor solubility as ferric oxide (Fe³⁺) particularly in highly oxygenated, neutral or alkaline soils (Hindt & Guerinot, 2022). This limited bioavailability often leads to iron deficiency in plants and in natural and agricultural environments iron fertilisation is often necessary (Zuo & Zhang, 2011). To counter this challenge plants have evolved specialised iron acquisition strategies. Dicots and non-grass monocots employ “strategy I” which relies on the reduction of iron prior to its import (Römheld, 1987; Eide *et al*., 1996). In the model strategy I plant *Arabidopsis thaliana* (*Arabidopsis*) this is conferred by the tripartite reductive import module consisting of H⁺-ATPase AHA2, FERRIC REDUCTION OXIDASE 2 (FRO2) and IRON-REGULATED TRANSPORTER 1 (IRT1)(Tsai & Schmidt, 2017). AHA2-mediated proton extrusion into the rhizosphere is thought to contribute to ferric iron (Fe³⁺) mobility through acidification of the rhizosphere (Santi & Schmidt, 2009). Acidification also lowers the redox potential to support Fe³⁺ reduction to ferrous iron (Fe²⁺) by FRO2, and consequently Fe^2+^ is taken up via IRT1 (Vert *et al*., 2002; Varotto *et al*., 2002). Graminaceous plants, on the other hand, utilise “strategy II,” marked by the release of mugineic acid-type phytosiderophores that chelate Fe³⁺ and form complexes that can be efficiently absorbed by specific root transporters (Takagi *et al*., 1984; Römheld & Marschner, 1986; Ishimaru *et al*., 2006; Murata *et al*., 2006; Nozoye *et al*., 2011)

In addition to acidification, several strategy I plants also exude specific iron-chelating coumarins to assist in iron solubilisation. Iron chelation is facilitated by catechol coumarins such as esculetin, fraxetin, and sideretin, with two hydroxyl groups in ortho positions (Hider *et al*., 2001; Mladěnka *et al*., 2010). The effectiveness of these coumarin metabolites is pH-dependent and expression of their biosynthesis genes responds to pH fluctuations (Sisó-Terraza *et al*., 2016; Tsai & Schmidt, 2017, 2020; Gautam *et al*., 2021; Paffrath *et al*., 2023). Plants also exude the non-catechol containing coumarin scopoletin which is not effective in mobilising iron (Schmid *et al*., 2014; Sisó-Terraza *et al*., 2016; Rajniak *et al*., 2018). The complexity of coumarin profiles and their biological activity is further increased by their ability to influence the composition and dynamics of the plant root microbiota. (Zamioudis *et al*., 2014; Stringlis *et al*., 2018; Voges *et al*., 2019; Harbort *et al*., 2020). Several coumarins, especially scopoletin, were reported to possess selective antimicrobial activities (Stringlis *et al*., 2019) and inhibit the growth of soil-borne fungi (Carpinella *et al*., 2005; Kai *et al*., 2006; Ba *et al*., 2017; Stringlis *et al*., 2018). Moreover, Harbort *et al* (2020) demonstrated that root commensal bacteria can alleviate iron starvation in *Arabidopsis* through interactions with root-secreted fraxetin in alkaline pH conditions. *Arabidopsis* mutants lacking coumarins harboured altered bacterial communities and had heightened iron stress sensitivity (Harbort *et al*., 2020).

The first step in the *Arabidopsis* coumarin biosynthesis pathway involves the 2-oxoglutarate-dependent dioxygenase FERULOYL-CoA 6’-HYDROXYLASE (F6’H1) (Kai *et al*., 2008; Schmidt *et al*., 2014; Schmid *et al*., 2014). F6’H1 converts feruloyl-CoA to 6-hydroxyferuloyl-CoA, which is then converted to scopoletin spontaneously or by the BAHD acyltransferase-COUMARIN SYNTHASE (COSY) in organs shielded from light (Kai *et al*., 2008; Vanholme *et al*., 2019). COSY was also found to produce esculetin *in vitro* from its hydroxycinnamoyl-CoA thioester (Vanholme *et al*., 2019). While the mode of esculetin biosynthesis in the plant remains unclear, Rajniak et al. (2018) suggested that F6’H1 is required, although *Arabidopsis* exudates contain relatively low levels of esculetin compared to scopoletin (Schmid *et al*., 2014; Paffrath *et al*., 2023). Fraxetin is synthesised from scopoletin by SCOPOLETIN-8-HYDROXYLASE (S8H) and can be further hydroxylated into sideretin by cytochrome P450 82C4 enzyme (CYP82C4) (Siwinska *et al*., 2018; Rajniak *et al*., 2018; Tsai *et al*., 2018). These last steps in the coumarin biosynthesis pathway (S8H and CYP82C4) are highly induced in response to iron deficiency and lead to higher coumarin exudation rates (Rodríguez-Celma *et al*., 2013; Fourcroy *et al*., 2014; Schmid *et al*., 2014; Sisó-Terraza *et al*., 2016; Tsai *et al*., 2018; Paffrath *et al*., 2023). Coumarins are secreted into the rhizosphere via ATP-binding cassette transporters like PLEIOTROPIC DRUG RESISTANCE 9 (PDR9) (Fourcroy *et al*., 2014; Ziegler *et al*., 2017), where they can directly contribute to iron uptake by chelating or reducing Fe^3+^ (Hider *et al*., 2001; Mladěnka *et al*., 2010; Schmidt *et al*., 2014; Paffrath *et al*., 2023).

While a critical role of coumarin interactions with root-associated bacterial microbiota for plant iron nutrition has been established (Harbort *et al*., 2020), contributions of fungal root endophytes and their interplay with root-secreted coumarins for plant iron nutrition remain unreported. Here we used a fungal endophyte, *Macrophomina phaseolina* MPI-SDFR-AT-0080 (F80), which is a member of the *Arabidopsis* mycobiota isolated from the roots of wild *Arabidopsis* populations and part of a microbial culture collection (Durán *et al*., 2018). We demonstrate that *M. phaseolina* F80 alleviates host iron-starvation in a coumarin-dependent manner. We uncover a previously undescribed mechanism by which a root endophytic fungus promotes plant iron nutrition through cooperation with the plant-derived coumarin scopoletin. Ex planta assays suggest that F80 mediates the conversion of scopoletin to the iron-mobilising catechol coumarin esculetin which then relieves plant iron deficiency by providing soluble Fe^3+^ to FRO2 for reduction and import via IRT1. Our data highlight a role of coumarin-microbiota interactions for plant iron nutrition across microbial kingdoms and further suggest physiological importance of the highly abundant coumarin scopoletin for plant iron nutrition in nature.

## Material and methods

### Growth conditions

The following *A. thaliana* genotypes were used: Columbia-0 wildtype (Col-0), *f6’h1-*1 AT3G13610; SALK_132418C (*f6’h1*), *s8h-1* AT3G12900; SM_3_27151 (*s8h*), *cyp82c4-1* AT4G31940; SALK_001585 (*cyp82c4*), *frd1-1* AT1G01580; N3777 (*fro2*), and *irt1-1* AT4G19690; SALK_024525 (*irt1*). *Arabidopsis* seeds were surface sterilised for 15min with 70% ethanol + 0.1% Tween-20, subsequently for 2min in 95% ethanol, and after drying, seeds were kept in sterile distilled water for 2-3 days in the dark at 4°C for stratification. For germination, seeds were placed onto square Petri plates containing ½× Murashige & Skoog (MS) medium (vitamins, 0.5 g/L MES, pH 5.7, 0.5% sucrose, 1% BD DIFCO™ Agar). The plates were sealed with micropore tape and placed vertically into a growth chamber (10 h light, 21°C; 14 h dark, 19°C) for 7 d.

### Fungal growth & hyphal harvesting

The fungus *Macrophomina phaseolina* (MPI-SDFR-AT-0080 v1.0) (F80) (Mesny *et al*., 2021), was grown on Potato Dextrose Agar (PDA) at 22°C for approx. 1 week. Hyphae were harvested with a sterilised scalpel, transferred to grinding tubes with metal beads, diluted to 100 mg hyphae/ml with 10 mM MgCl₂, and ground for 5 min. To obtain heat-killed (HK) fungus, samples were incubated at 99°C for 1 h.

### Gnotobiotic agar-plate system

The gnotobiotic agar-based system was adapted from (Harbort *et al*., 2020) - STAR protocols. Adjusted half strength MS media without iron (750 µM MgSO_4_, 625 µM KH_2_PO_4_, 10.3 mM NH_4_NO_3_, 9.4 mM KNO_3_, 1.5 mM CaCl_2_, 55 nM CoCl_2_, 53 nM CuCl_2_, 50 µM H_3_BO_3_, 2.5 µM KI, 50 µM MnCl_2_, 520 nM Na_2_MoO_4_, 15 µM ZnCl_2_, and 9.4 mM KCl, with 1% BD DIFCO™ Agar, Bacteriological) was prepared from stock solutions and supplied with 10 mM MES, 50 μM iron (FeEDTA or FeCl₃), and fungal hyphae (1:2000 dilution, 50 μg/ml final concentration) after autoclaving. For coumarin supplementation experiments, coumarin were added from 100 mM DMSO stocks to a final concentration of 10 μM or an equal amount of DMSO as control. After solidifying, the top approx. 2.5 cm agar was removed from 12x12 square petri plates filled with 50 ml of medium, and seven germinated seedlings were transferred per plate (four replicate plates per experiment). Plates were sealed with micropore tape and vertically grown for 2 weeks in a growth chamber (10 h light, 21°C; 14 h dark, 19°C) with roots shielded from light. After 2 weeks, shoot samples were collected for chlorophyll measurements and whole shoots and roots were harvested to record shoot fresh weight (SFW) and assess fungal colonisation, respectively.

### Shoot chlorophyll analysis

The protocol for chlorophyll extraction and quantification was based on Harbort et al., (2020) and Hiscox & Israelstam, (2011). 5-7 shoots from each plate were pooled, weighed and kept at −80°C until processing. To each sample, 1 ml DMSO per 30 mg of leaf tissue was added and they were incubated for 45 min at 65 °C with 300 rpm agitation. Chlorophyll absorbance was measured at 652 nm on a Nanodrop spectrophotometer (Thermo Fisher Scientific).

### Shoot mineral element analysis

Mineral element analysis of whole shoots was performed as described previously (Paffrath *et al*., 2023) Shoot samples were dried to constant weight at 65°C and weighed into polytetrafluoroethylene tubes. Dried plant material was digested using concentrated HNO₃ (67–69% v/v, Bernd Kraft) and subjected to pressurised digestion in a high-performance microwave reactor (UltraCLAVE IV, MLS GmbH). After digestion, the samples were diluted with deionised water (Milli-Q Reference A+, Merck Millipore) and analysed using high-resolution inductively coupled plasma mass spectrometry (HR-ICP-MS) (ELEMENT 2, Thermo Scientific, Germany). Element standards were prepared from certified reference single standards from CPI-International (USA).

### Fungal colonisation assays

To assess fungal colonisation of *Arabidopsis* roots at 14 days post-inoculation (dpi), roots were collected into Lysing Matrix E 2 ml tubes (MP Biomedicals, USA). The tube content was crushed for 30 s at 6200 rpm. 980 μl sodium phosphate buffer (MP Biomedicals, USA) and 122.5 μl MT-buffer (MP Biomedicals, USA) were added, followed by two additional 30-secrounds of crushing at 6200 rpm. The samples were then centrifuged for 15 min at 13.000 rpm. 50 μl binding matrix (MP Biomedicals, USA) per sample was added to a 96-well plate. A filter plate (Acroprep Advance, 0.2 μm Supor filter, Pall) was placed on top of the 96-well plate, and 150 μl supernatant from each sample was transferred onto the filter plate. The plate was centrifuged for 20 min at 1500 rpm, thereafter the flow-through was mixed with the binding matrix, by shaking for 3 min. 190 μl of each sample was transferred to a second filter plate and washed twice with 150 μl SEWS-M (MP Biomedicals, USA) by centrifuging for 5 min at 1500 rpm and the centrifuging step was repeated to dry the samples. DNA was eluted with 30 μl of ddH2O by centrifuging at 1500 rpm for 5 min, and stored at −20 °C until qPCR analysis. Fungal colonisation was then measured by qPCR as described (Mesny et al., 2021). Briefly, ITS1F/ITS2 primers for fungal ITS1 and UBQ10F/UBQ10R primers for *Arabidopsis* Ubiquitin10 were used. Reactions included iQ™ SYBR® Green Supermix, primers, and 1 µl DNA template. qPCR was run on a BioRad CFX Connect with 95°C denaturation (3 min), followed by 39 cycles of 95°C (15 s), 60°C (30 s), and 72°C (30 s). Colonisation index was calculated as 2^-Cq(ITS1)/Cq(UBQ10)^.

### 96-well fungal culture assay

Culture medium containing half strength MS medium without Fe, ARE and vitamins (Table S1) was prepared, the pH was adjusted to pH 5.7 (buffered with 10 mM MES) with KOH and sterile-filtered (0.22 μm) using vacuum filtration. The medium was supplemented with scopoletin (0, 0.0625, 0.25, 1, or 2 mM) or pimaricin (0.1 mg/ml final concentration, used as an antifungal growth control) and 190µl was added per well in a 96-well plate. The wells were inoculated with 10 μl of fungal hyphal solution (stock solution of 100 mg mycelium per ml in 10mM MgCl_2_; final concentration 5mg/ml) 10 μl of 10mM MgCl_2_ as a blank control. The plates were sealed and incubated for 12 days at 22 °C with shaking (140 rpm), OD600 and scopoletin fluorescence (excitation 385 nm, emission 470 nm) measurements were taken daily around the same time using a TECAN spectrophotometer (Infinite® M Plex, Tecan Trading AG, Switzerland). To ensure consistency across experiments, detected fluorescence was normalised to OD600 and the initial fluorescence at day 0. To evaluate potential fungal growth inhibition by scopoletin, daily OD600 values were normalised to growth in the absence of scopoletin.

### Metabolite profiling

For F80 liquid culture time-course assays, pure ethyl acetate (Honeywell Riedel-de HaënTM, LC-MS grade) was used to extract metabolites from fungal culture pellet and supernatant fractions. Extraction was performed by adding 500 µl of ethyl acetate to 1.5 ml fungal culture fractions, followed by 30 s vortexing and 10 min sonic bath. The sample was centrifuged for 10 min (13000 rpm, 4°C) and 450 µl of upper phase was collected in a new 1.5 ml test tube. The extraction was repeated two times. The combined solvent extract was dried in a TurboVap^®^ Classic LV (Biotage) under nitrogen flow, and samples were stored at −80°C.

For exudate profiling in the plant medium, 5 ml pure ethyl acetate (Honeywell Riedel-de HaënTM, LC-MS grade) was added to 50 ml plant agar medium (harvested 1 week after transfer of 9-day-old seedlings), and samples were vortexed for 30 s and centrifuged for 10 min (4°C, 4000 rpm). The supernatant was collected and two more rounds of extraction were performed. Combined extracts were dried as described above and samples stored at −80°C.

For coumarin profiling of plant roots (harvested 1 week after transfer of 9-day-old seedlings on half strength MS medium), 200 mg (FW) ground root samples were extracted using a modified version of Blight & Dyer extraction. First, 900 µl dichloromethane/ethanol 2:1 and 100 µl HCl (pH 1.4) were consecutively added to each sample, followed by 30 s vortexing and 5 min centrifugation (4°C, 13600 rpm). The upper phase was discarded and the lower phase (approx. 700 µl) was collected into 2 ml test tubes. After adding 500 µl tetrahydrofuran to the remaining pellets, samples were vortexed for 30 s and centrifuged for 5 min (4°C, 13600 rpm). The supernatant was collected and combined with the lower phase fraction. The solvent mixture was dried as described above, and the samples stored at −80°C.

Before measurement, each sample was resuspended in 100 µl of pure methanol (Honeywell Riedel-de HaënTM, LC-MS grade) and transferred to a liquid chromatography vial with insert. Separation of metabolites by ultrahigh-performance liquid chromatography (UHPLC) was performed on a Nucleoshell RP18 (2.1 × 150 mm, particle size 2.1 μm, Macherey & Nagel, GmbH, Düren, Germany) using an ACQUITY UPLC System, equipped with a Binary Solvent Manager and Sample Manager (20 μl sample loop, partial loop injection mode, 2 μl injection volume, Waters GmbH Eschborn, Germany). Eluents A and B were aqueous 0.3 mmol/L NH4HCOO (adjusted to pH 3.5 with formic acid) and acetonitrile, respectively. Elution was performed isocratically for 2 min at 5% eluent B, from 2 to 7 min with a linear gradient to 30% B, from 7-9 min at 95% B, then 1 min at 95% B, from 10 to 11 min at 5% B, and then another minute at 5% B. The flow rate was set to 400 μl min−1 and the column temperature was maintained at 40 °C. Metabolites were detected by negative electrospray ionisation and mass spectrometry (ESI-MS). Serial dilutions of standards of scopoletin, esculetin, fraxetin and 3- (2,4-dihidroxy-5-methoxyphenyl) propanoic acid were injected every 24 samples for identification and quantification purposes. Mass spectrometric analysis was performed by MS-QTOF-IDA-MS/MS (ZenoTOF 7600, AB Sciex GmbH, Darmstadt, Germany) and controlled by SCIEX OS software (AB Sciex GmbH, Darmstadt, Germany). The source operation parameters were as follows: ion spray voltage, −4500 V; ion source gas 1, 60 psi; source temperature 600 °C; ion source gas 2, 70 psi; curtain gas, 35 psi. Instrument tuning and internal mass calibration were performed every 5 samples with the calibrant delivery system applying X500 ESI Calibrant Solution in negative mode tuning (AB Sciex GmbH, Darmstadt, Germany). Data analysis was performed in SCIEX OS software, and MS-DIAL software (Tsugawa *et al*., 2015).

### Ferrozine iron mobilisation assay

To remove medium residues from the hyphal solution, fungal hyphae (prepared as described above) were washed three times with MgCl_2_ by centrifugation (5 min at 11,000 xg), and resuspended in fresh MgCl_2_. Coumarins were supplemented to the culture medium (half strength MS medium without iron, ARE and vitamins (Table S1), 10 mM MES, pH 5.7) to a final concentration of 400 μM. Cultures were inoculated with washed hyphae and grown as described for the 96-well fungal culture assay (22°C, 140 rpm, dark). Samples were collected at 0, 6, and 9 dpi, centrifuged at 3800 rpm for 20 min, the supernatant was transferred into a new 96 well plate and stored at −80 °C until processing.

To assess iron mobilisation, 120 μl sample was transferred into a 96-well plate and supplemented with a final concentration of 0.1 mM FeCl₃ and 0.4 mM ferrozine. Absorbance at 562 nm was measured using a TECAN spectrophotometer (Infinite® M Plex, Tecan Trading AG, Switzerland) before and after ferrozine addition. Controls included wells with water, 0.1 mM FeCl₃ (negative control), and 0.1 mM FeEDTA (positive control). To quantify total mobilised iron, immobile iron was removed by centrifugation (3800 rpm, 15 min), and the supernatant was transferred to a new 96-well plate. Samples were then incubated with 10 mM HCl and 10 mM ascorbic acid for 20 min in darkness to reduce the mobile iron fraction, followed by absorbance measurement at 562 nm. All values were normalised to FeEDTA samples (before reduction), and FeSO_4_ calibration curves were used for quantification of mobile iron.

## Results

### *M. phaseolina* (F80) enhances iron nutrition in *Arabidopsis* under iron limiting conditions

To test whether fungal endophytes can improve plant iron nutrition, we modified a previously reported gnotobiotic agar-based plant growth assay that allows precise modulation of iron bio- availability and the co-cultivation of *Arabidopsis* plants with root microbiota members (Harbort *et al*., 2020 - STAR protocols). The system relies on half strength MS medium buffered at pH 5.7 with MES and supplemented with either unchelated iron (FeCl_3_) or chelated iron (FeEDTA), which reflect bio-unavailable iron (unavFe) and bio-available iron (avFe) conditions, respectively. Using this system, we assessed the ability of the fungal root endophyte F80 to enhance the performance of *Arabidopsis* Col-0 wild-type plants under unavFe conditions. F80 was identified previously as part of a culture collection of 41 fungal isolates representative of the *Arabidopsis* root mycobiota (Mesny *et al*., 2021). Plant performance was evaluated after two weeks of co-cultivation with fungus or mock treatments using shoot fresh weight (SFW) and shoot total chlorophyll concentration as a proxy for iron status. Three-week-old *Arabidopsis* seedlings exhibited enhanced growth and reduced leaf chlorosis when inoculated with F80 under iron-limiting conditions, closely resembling the avFe phenotype (Fig. 1a). Plants grown at unavFe and inoculated with F80 exhibited a significant increase in SFW and shoot chlorophyll concentration compared to mock-treated controls (Fig. 1b,c). By contrast, we did not observe F80-mediated plant growth promotion or increased chlorophyll under avFe conditions (Fig. 1b,c). Plants inoculated with heat-killed F80 behaved similarly to mock-treated plants (Fig. S1), highlighting the importance of live fungus and that putative inoculum-derived iron contamination is negligible. To ascertain whether the observed phenotypes reflect iron status of the plant, the shoot elemental content was measured by HR-ICP-MS (Fig. 1d). This analysis showed that F80-inoculated plants accumulated significantly more iron in their shoots under unavFe compared to mock conditions, almost reaching the iron content of plants grown at avFe (Fig. 1d). In line with SFW and shoot chlorophyll concentration, no F80-mediated increase in shoot iron content was observed in avFe conditions (Fig. 1d). Taken together, these results show that F80 mediates *Arabidopsis* iron nutrition specifically in conditions of limited iron bioavailability. This implies that F80 improves plant access to immobile iron pools in the growth medium.

**Fig. 1.**
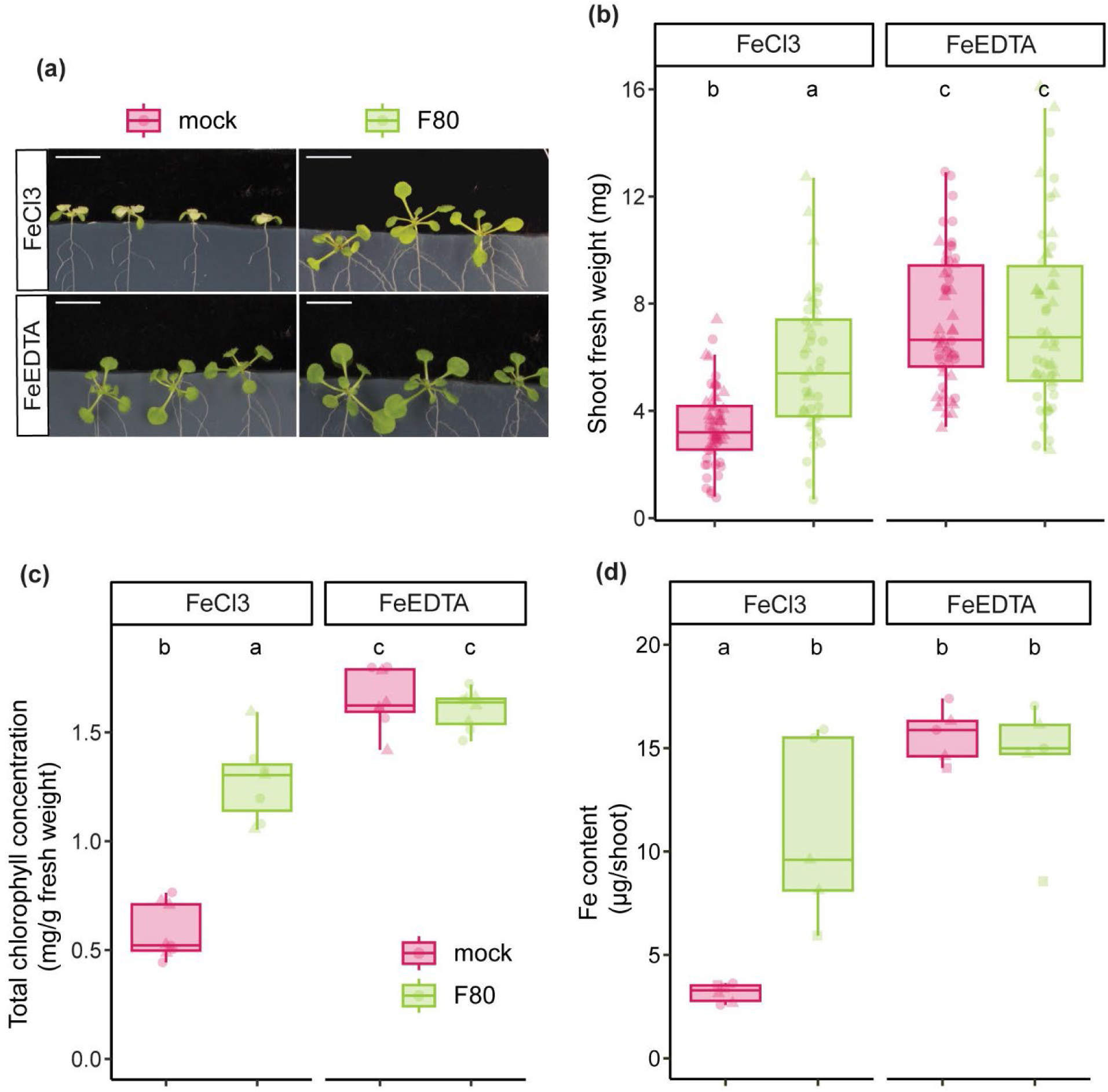
Fungal endophyte *M. phaseolina* (F80) improves *Arabidopsis* iron nutrition under iron limitation. (a) representative phenotypes, (b) SFW, (c) shoot chlorophyll concentration, and (d) shoot Fe content of Col-0 *Arabidopsis* seedlings. 7-day-old seedlings were transferred to half-strength MS medium and grown for 2 weeks on mock or F80 inoculated plates in unavFe (50µM FeCl3) or avFe (50µM FeEDTA) conditions. Letters indicate significant pairwise differences between groups (*p*-adj≤0.05) by Dunn pairwise comparison test with Benjamini-Hochberg correction (b,c) or a Tukey’s HSD corrected for multiple comparisons (d). Scale bars, 1 cm (a). Data are from two (b,c) or three (d) full factorial replicates (represented by different shapes).

### Root-derived scopoletin and host iron-reductive import machinery are essential for F80- mediated plant iron nutrition

Given that F80-mediated plant iron nutrition is specific to iron-limiting conditions and the established link between iron limitation-induced coumarin production and bacterial-mediated improvements in plant iron nutrition (Harbort *et al*., 2020), we investigated whether F80- mediated improved in iron nutrition of *Arabidopsis* also requires plant-derived coumarins. To this end, we grew the coumarin-deficient *Arabidopsis* mutant *f6’h1* lacking the first step of the coumarin biosynthesis pathway (Kai *et al*., 2008; Schmidt *et al*., 2014; Schmid *et al*., 2014) in unavFe conditions. Compared to Col-0, *f6’h1* plants exhibited reduced ability to tolerate iron deficiency, indicated by their lower SFW and shoot chlorophyll concentration under mock conditions (Fig. 2a,b). While F80-mediated rescue of plant iron deficiency was observed in Col- 0 plants, F80 failed to improve the performance of *f6’h1* plants, with both SFW and chlorophyll levels remaining below the mean values observed for Col-0 (Fig. 2a,b). Consistently, shoot iron content in *f6’h1* plants did not significantly increase upon F80 inoculation and remained comparable to mock-treated controls (Fig. 2c).

**Fig. 2.**
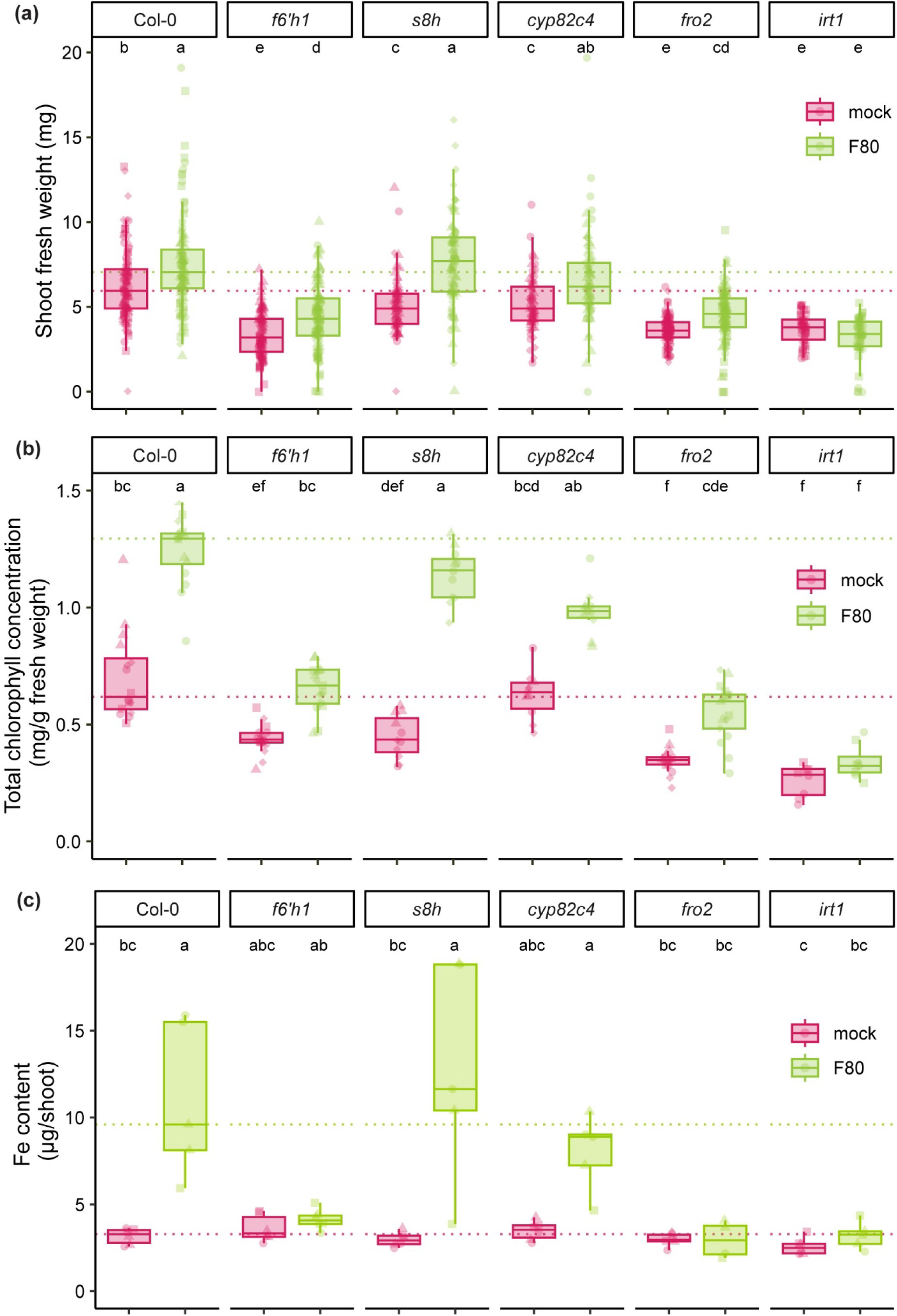
Rescue of iron deficient *Arabidopsis* plants by the fungal root endophyte F80 requires the coumarin scopoletin and the plant’s iron reductive import system. (a) SFW, (b) shoot chlorophyll concentration and (c) shoot Fe content measured by ICP-MS at 2 weeks of growth after transfer of the indicated mutants in the coumarin biosynthesis pathway and the iron reductive import mechanism. 7-day-old seedlings were transferred to half strength MS medium with unavFe (50µM FeCl3) at pH 5.7 mock or inoculated with F80. Dashed line indicates the mean of Col-0. Letters indicate significant pairwise differences between groups (*p*-adj≤0.05) by a Dunn pairwise comparison test with Benjamini-Hochberg correction. Statistical significance within genotypes was determined by a Kruskal-Wallis test. Data are from four (a,b) or three (c) full factorial replicates (represented by different shapes).

To examine whether F80 interacts with specific coumarins to facilitate rescue of plant iron deficiency, we used the *s8h* and *cyp82c4* mutant lines which fail to produce fraxetin and sideretin, respectively, yet retain the ability to synthesise scopoletin (Rajniak *et al*., 2018; Tsai *et al*., 2018). Inoculation of *s8h* plants with F80 resulted in a significant increase in SFW and shoot chlorophyll concentration, reaching levels comparable to those of F80-inoculated Col-0 plants. This trend was also observed for shoot iron content, as *s8h* plants treated with F80 accumulated similar levels of iron as Col-0 (Fig. 2c). Similarly, F80 inoculation improved the performance of *cyp82c4* plants reflected in increased SFW, chlorophyll concentration and shoot iron content compared to mock-treated controls, although levels in *cyp82c4* remained slightly lower than those of Col-0 and *s8h* plants (Fig. 2a,b,c). These data show that the fungal root endophyte alleviates plant iron starvation in a coumarin-dependent manner and that the coumarin scopoletin interacts with F80 to rescue host iron deficiency.

To investigate whether F80-mediated plant iron rescue depends on the plant’s ability to reduce ferric (Fe^3+^) and import ferrous (Fe^2+^) iron, we assessed mutants of *FRO2* and *IRT1*, two key components of the iron acquisition system in *Arabidopsis*. As previously reported, the *fro2* and *irt1* mutants were acutely sensitive to iron limitation, resulting in small and chlorotic plants under unavailable iron conditions (Robinson *et al*., 1999; Vert *et al*., 2002; Varotto *et al*., 2002; Paffrath *et al*., 2023). F80 failed to effectively increase *fro2* and *irt1* SFW, shoot chlorophyll concentration and shoot iron content (Fig. 2a,b,c). This suggests that the F80-mediated rescue involves ferric iron mobilisation and requires downstream Fe^3+^ reduction by FRO2 as well as ferrous iron transport by IRT1. The concentration of other elements in the shoots were also evaluated for all genotypes by HR-ICP-MS, however no similar trends were observed (Fig. S5). The observed differences in F80-mediated plant iron nutrition were not attributed to variation in fungal colonisation, as F80 colonisation levels were similar in all plant genotypes (Fig. S2c). Analysis of all plant genotypes at avFe confirmed that F80-mediated rescue is specific to iron deficiency as there were no differences observed in shoot chlorophyll of mock versus F80 inoculated plants (Fig. S2a,b). Additionally, presence of FeEDTA completely restored plant growth of all tested coumarin mutants, indicating that coumarins are not required for plant growth at avFe. As anticipated, FeEDTA did not fully restore chlorophyll levels in *fro2* and *irt1*, as iron reduction and uptake are still required under this condition (Fig. S2a,b). For all genotypes except *s8h*, an increase in SFW was observed in avFe conditions in the presence of F80, which might reflect a general beneficial effect of the fungus on plant growth. The results suggest that F80-mediated *Arabidopsis* iron acquisition depends on plant coumarin synthesis, especially scopoletin, and requires a functional plant reductive iron transport machinery to facilitate iron uptake.

### F80 converts scopoletin into iron-chelating esculetin

Scopoletin is recognised for its selective antimicrobial activity (Sun *et al*., 2014; Zamioudis *et al*., 2014; Lemos *et al*., 2020; Harbort *et al*., 2020)). To evaluate the sensitivity of F80 to scopoletin, an *in vitro* time-course assay was performed to monitor the fungal growth in the presence of 2 mM scopoletin in a liquid culture. To additionally assess potential F80-mediated alterations of scopoletin, we tracked scopoletin-derived fluorescence in the cultures over time under long-wave UV (Goodwin & Kavanagh, 1949). Under mock conditions, the expected scopoletin-derived fluorescence remained largely constant, confirming stability of scopoletin in the absence of the fungus. This is consistent with a previous study using mass spectrometry-based quantification of coumarins (Rajniak *et al*., 2018). In the presence of F80, scopoletin-derived fluorescence decreased rapidly between 3- and 6-days post-inoculation (dpi) (Fig. 3a). A scopoletin alteration effect was also indicated by the fungal growth pattern: relative growth of F80 (growth in 2 mM scopoletin/growth in control) decreased from 2 to 5 dpi but recovered from 6 dpi onwards, coinciding with a near-complete disappearance of scopoletin fluorescence (Fig. S3b). These data suggest that F80 can evade scopoletin antimicrobial activity by a possible detoxification mechanism that alters scopoletin.

**Fig. 3.**
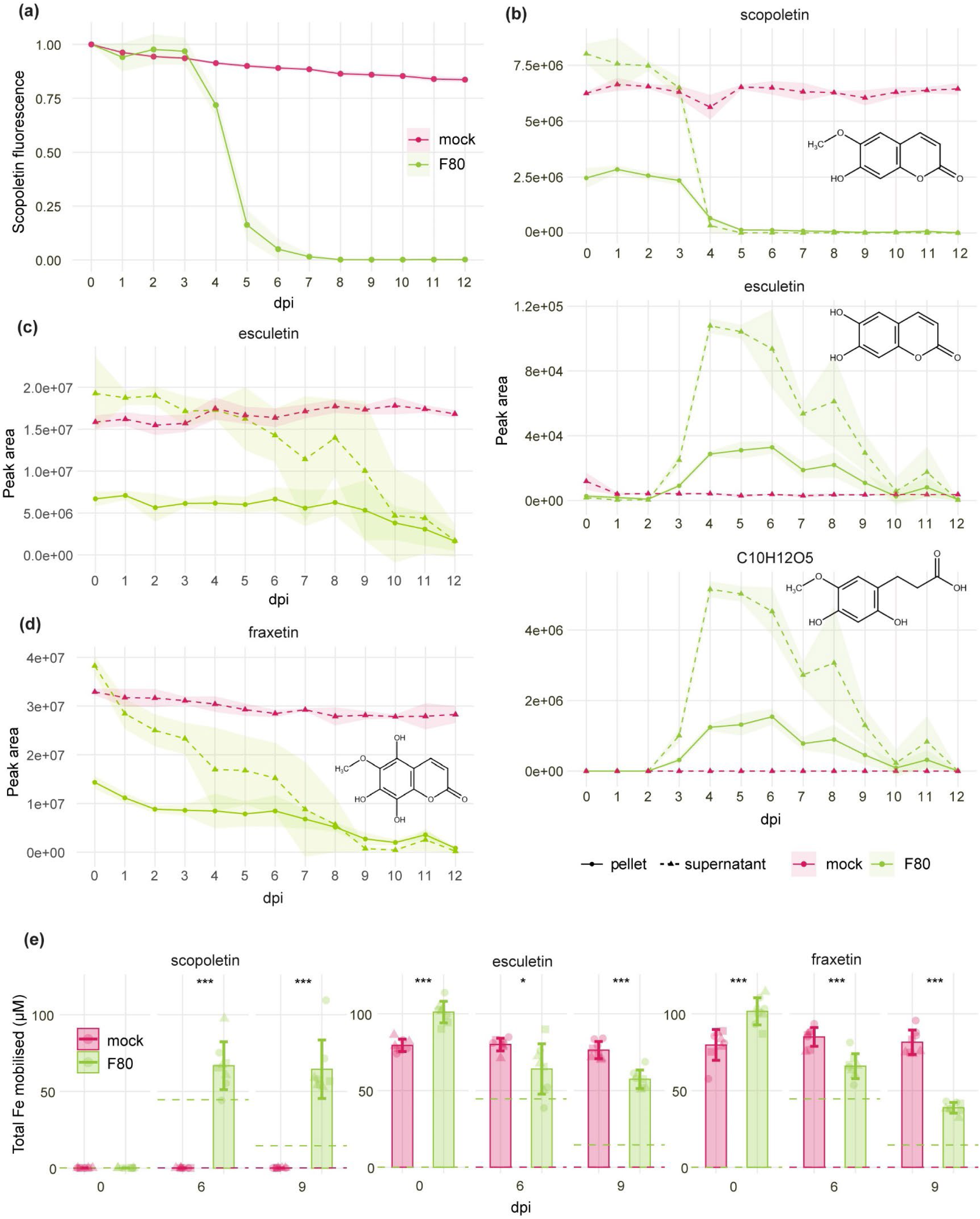
The fungal endophyte F80 converts scopoletin into esculetin and further degradation compounds. (a) Time course of 0-12 dpi displaying specific scopoletin fluorescence measured in *in vitro* mock and F80 cultures supplemented with 2mM scopoletin (excitation at 385nm and emission detected at 470nm). Data combined from 3 biological replicates. The curve-shade indicates the standard error. The Area Under the Curve of graph (a) is shown in Fig. S3a. (b) MS-QTOF-IDA-MS/MS peak area of scopoletin, esculetin and 3-(2,4-dihydroxy-5-methoxyphenyl) propanoic acid (C10H12O5) in F80 culture supernatant (dashed line) and the fungal pellet (full line) grown for 0-12 dpi supplemented with 2mM scopoletin. Curve-shade indicates mean ± SD. Absolute concentration values are presented in Figure S3d. (c) MS-TOF-IDA-MS/MS peak area of esculetin in F80 culture supernatant (dashed line) and the fungal pellet (full line) grown for 0-12 dpi supplemented with 2mM esculetin. Curve-shade indicates mean ± SD. Absolute concentration values are presented in Figure S3e. (d) MS-TOF-IDA-MS/MS peak area of fraxetin in F80 culture supernatant (dashed line) and the fungal pellet (full line) grown for 0-12 dpi supplemented with 2mM fraxetin. Curve-shade indicates mean ± SD. Absolute concentration values are presented in Figure S3f. (e) Total amount of iron mobilised from poorly soluble FeCl₃ (0.1 mM) in the F80 culture supernatant or axenic medium (mock) supplemented with scopoletin, fraxetin, or esculetin (final concentration of 0.4 mM) at 0, 6, or 9 dpi. Mobilised iron was assessed spectrophotometrically based on a Fe(II)-ferrozine assay. Control values for treatments without fungus (red) or without coumarins (green) are included as dashed lines. Error bars represent mean ± SD, statistical differences between treatments were assessed using *t*-tests for normally distributed residuals or, otherwise, Wilcoxon rank-sum tests. Significance levels are indicated by asterisks with \**p*<0.05, \*\**p*<0.01, and \*\*\**p*<0.001. All panels represent data from three biological replicates (represented by different shapes).

To validate whether F80 alters scopoletin, liquid cultures of the fungus were separated into supernatant and mycelia, and metabolites were profiled by UHPLC-MS/MS. The MS/MS data corroborated the fluorescence-based results in showing a steep decrease in scopoletin levels from 3 to 6 dpi with F80 (Figs 3b, S3d). Notably, a different coumarin molecule, esculetin, was detected in the samples supplemented with scopoletin, and displayed increased abundance from 3 dpi onwards. At the same time, additional degradation products containing hydrolysed lactone rings were observed. Untargeted UHPLC-MS/MS profiling of the samples revealed that the most abundant compound detected, besides scopoletin, was 3-(2,4-dihydroxy-5- methoxyphenyl) propanoic acid (C10H12O5) (Figs 3b, S3d). At 12 dpi, all coumarin compounds, including scopoletin, esculetin, and 3-(2,4-dihydroxy-5-methoxyphenyl) propanoic acid, were nearly undetectable. This suggests that further modification or complete degradation through F80, or compound instability, occurs in this assay. Both, esculetin and the open-lactone structure were detected exclusively in scopoletin-supplemented samples in the presence of the fungus but not in fungal cultures alone (Figs 3b, S3d).

Given that esculetin, an iron-mobilising coumarin (Schmid *et al*., 2014; Paffrath *et al*., 2023), was identified as a scopoletin-conversion product, we next tested the interaction of esculetin with F80. First, esculetin stability in F80 cultures supplemented with 2 mM esculetin was assessed by measuring its levels in the supernatant and fungal pellet over time. Esculetin showed only a slow decline and was still detectable at 12 dpi (Figs 3c, S3e). In parallel, since fraxetin accumulates in the roots of *cyp82c4* plants which support F80-mediated rescue, the stability of fraxetin was evaluated in the presence of F80 in culture (Fig. 2a,b,c). Compared to esculetin, fraxetin was more rapidly depleted by the fungus, especially in the supernatant (Figs 3d, S3f). To further investigate the interaction of F80 with these coumarins, relative growth of F80 in the presence of esculetin or fraxetin was evaluated over time. In comparison to F80 cultures supplemented with scopoletin, F80 cultures supplemented with esculetin or fraxetin did not show a decrease in relative growth (Figs S3b,c). The observed F80 conversion of scopoletin to esculetin, a compound that does not affect F80 growth, further supports a fungal detoxification mechanism.

Because our *in planta* data imply a role for F80 in improving plant access to immobile ferric iron through interactions with scopoletin - a coumarin which lacks a catechol moiety and *in vitro* iron mobilising capacity (Fig. 2a,b,c) (Schmidt *et al*., 2014; Rajniak *et al*., 2018; Paffrath *et al*., 2023) - we hypothesised that F80-mediated alteration of scopoletin is the cause of iron mobilisation to the plant. To test this, we made use of a spectrophotometric assay which quantifies mobile iron based on the ferrozine-Fe(II) complex. Fraxetin and esculetin were included in the analysis because esculetin was detected in the F80-scopoletin cultures (Figs 3b, S3d) and fraxetin produced in F80-rescued *cyp82c4* plants might also interact with the fungus. As expected, the catecholic coumarins fraxetin and esculetin, but not scopoletin, mobilised iron under mock conditions (Fig. 3e). However, when F80 was cultured with these coumarins, differential effects on total iron mobilisation were observed. In F80 cultures supplemented with scopoletin, there was an increase in mobilised iron at 6 and 9 dpi (Fig. 3e), perhaps due to the conversion of scopoletin to esculetin by F80 (Fig. 3b). Conversely, total iron mobilisation in cultures supplemented with fraxetin or esculetin in the presence of the fungus decreased significantly at 6 and especially 9 dpi (Fig. 3e). The results suggest that F80 affects iron mobilisation differently depending on the coumarin substrate it encounters. While scopoletin likely serves as a precursor to iron-chelating compounds such as esculetin, fraxetin and esculetin participate directly in iron-binding dynamics.

These findings show that an interaction between F80 and scopoletin yields synergistic amounts of mobilised iron. Put together with F80’s capability to convert scopoletin to the catechol coumarin esculetin, we reasoned that increased iron mobility might be due to the presence of esculetin. This could explain the observed F80-mediated *Arabidopsis* iron nutrition.

### F80 and scopoletin cooperate to mobilise ferric iron in the rhizosphere

To assess whether F80 interaction with coumarins observed in culture translate into iron mobilisation effects *in planta*, we conducted coumarin supplementation assays using the coumarin-deficient *f6’h1* mutant. This approach provided a coumarin-free plant background, enabling evaluation of the importance and sufficiency of specific coumarins for plant physiological fitness and growth under iron limitation. Based on the positive interaction of F80 and scopoletin in enhancing iron mobilisation and growth recovery of *f6’h1* plants at avFe (Fig. S2a,b), we hypothesised that combinatorial treatment of the plant growth medium with F80 and scopoletin would result in increased mobile ferric iron and, in turn, restore iron limiting plant growth of *f6’h1*. Addition of 10 µM scopoletin to the plant growth medium under mock (non-F80) conditions did not significantly increase SFW or chlorophyll concentration in *f6’h1* to levels observed in Col-0 inoculated with F80 (Fig. 4a,b). When combined with F80 inoculation, scopoletin supplementation restored F80-mediated rescue, with both SFW, chlorophyll and shoot iron concentrations being similar to Col-0 inoculated with F80 (Figs. 4a,b, S5). Strikingly, the cooperative effect of F80 and scopoletin resembled the mock phenotypes of 10 µM fraxetin or esculetin supplementations, which increased SFW and chlorophyll concentration to levels comparable to Col-0 with F80 (Fig 4a,b). Supplementation of fraxetin or esculetin in the presence of F80 resulted in slightly reduced SFW and chlorophyll levels compared to the corresponding mock conditions (Fig. 4a,b), in line with the fungal culture supernatant iron mobilisation data (Fig. 3e).

**Fig. 4.**
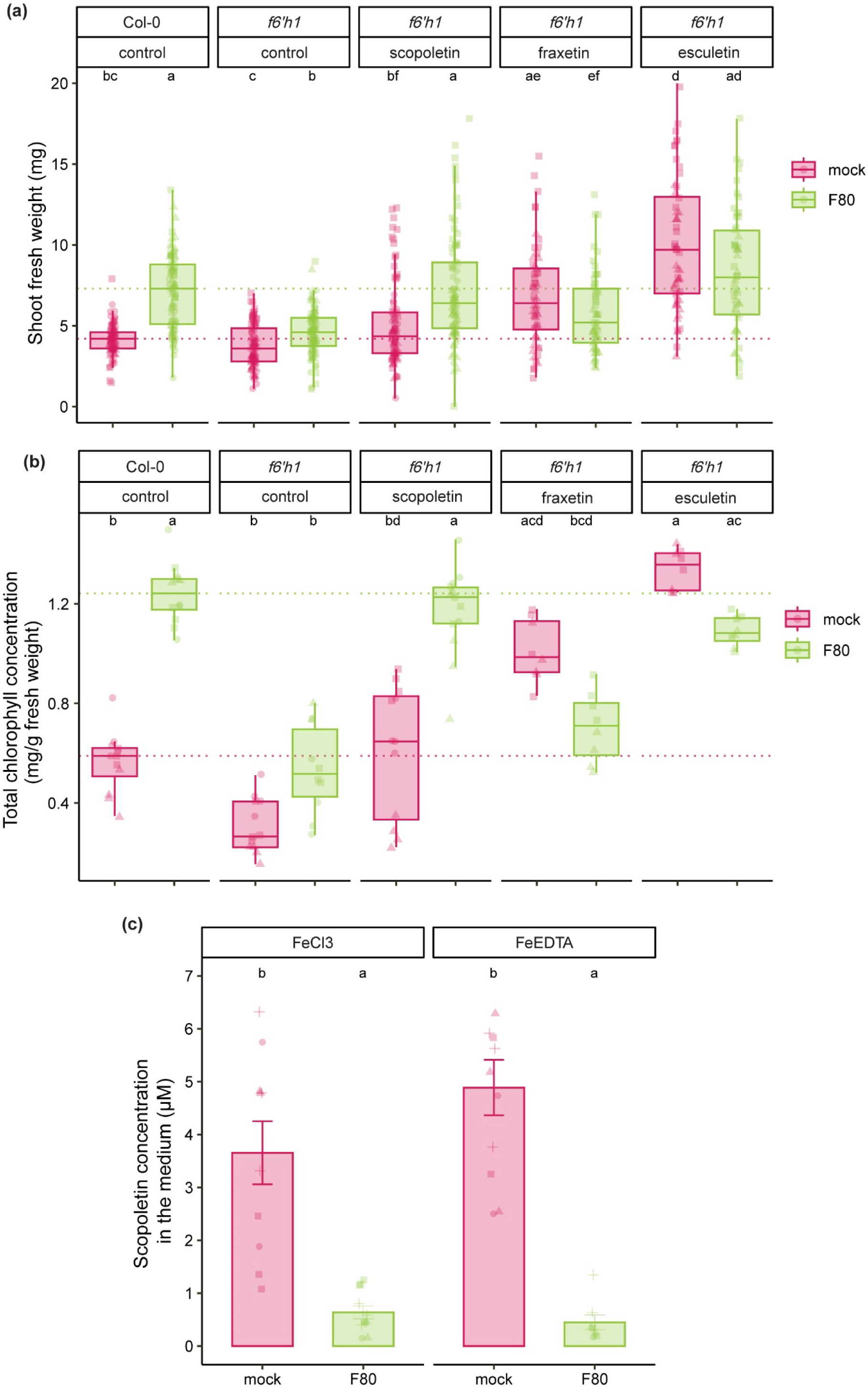
Interaction between the fungal endophyte F80 and scopoletin rescues the coumarin-deficient mutant *f6’h1*. (a) SFW and (b) shoot chlorophyll content of Col-0 and *f6’h1* plants 2 weeks past inoculation. 7-day-old seedlings were transferred to half-strength MS medium with unavailable iron (50 µM FeCl3) at pH 5.7, supplemented with 10 µM of the indicated coumarins or an equal amount of DMSO for the control, and mock or inoculated with F80. Dashed line indicates the mean of Col-0. Letters indicate significant pairwise differences between groups (*p*-adj≤0.05) by a Dunn pairwise comparison test with Benjamini-Hochberg correction. Data are from three full factorial replicates (represented by different shapes). (c) The concentration of scopoletin exuded after 1 week in the medium of 9-day-old seedlings transferred to half-strength MS medium with unavailable (50 µM FeCl3) or available (50 µM FeEDTA) iron at pH 5.7 mock or inoculated with F80. Metabolites were extracted from approximately 50 ml of the medium and analysed with MS-TOF-IDA-MS/MS. Bars represent the mean from 4 biological replicates (n=2-3) with standard error bars. Letters indicate significant pairwise differences between groups (*p*-adj≤0.05) by a Tukey’s HSD corrected for multiple comparisons.

To further understand the role of F80 in the iron import pathway of *Arabidopsis* plants, scopoletin, fraxetin or esculetin were added to the growth medium of *fro2* mutant plants. Importantly, none of these compounds significantly enhanced SFW or chlorophyll concentration in *fro2*, either alone or in combination with F80 (Fig. S4a,b). This underscores the necessity of a functional FRO2 for F80-mediated alleviation of plant iron deficiency. It also supports the notion that F80 cooperatively interacts with scopoletin to mediate ferric iron mobilisation in the rhizosphere. To critically test the impact of chemical iron mobilisation on *f6’h1* and *fro2* mutants, 10 µM NaEDTA (a synthetic iron chelator) was added to the medium. In *f6’h1* plants, NaEDTA supplementation almost fully rescued the iron-deficiency phenotype under mock and F80-inoculated conditions (Fig. S4c,d). This aligns with a previous study showing that iron chelators like NaEDTA can bypass the requirement for microbial interactions by artificially increasing iron solubility (Jin *et al*., 2006). By contrast, no significant NaEDTA- mediated rescue was observed in *fro2* mutants (Fig. S4c,d). This result reinforces the conclusion that FRO2 is essential for the iron uptake pathway when bio-available ferric iron is present in the form of Fe(III)-chelates (Figs. S4c, S3d). F80 colonisation of roots remained consistent across all experimental conditions, indicating that differences in rescue phenotypes were not due to variation in fungal colonisation (Fig. S4e).

To test whether F80 can also use plant-synthesised scopoletin, we quantified differences in scopoletin concentration in root exudates of Col-0 plants grown over 7 d in the presence or absence of F80. HRLC-MS/MS analysis revealed that the presence of F80 strongly decreased scopoletin levels in the agar medium in both unavFe and avFe conditions (Fig. 4c). Levels of scopolin and scopoletin quantified inside root tissues from this experiment revealed no significant differences between treatments (Fig. S4f). Altogether, the data suggest that F80 and scopoletin interact cooperatively in the rhizosphere to mediate ferric iron mobilisation upstream of the FRO2-IRT1 reductive import module.

## Discussion

Coumarins are crucial for plant iron uptake in conditions of low iron availability. Scopoletin is one of the most abundant coumarins in root exudates of *Arabidopsis* plants grown under iron-limiting conditions (Sisó-Terraza *et al*., 2016; Rosenkranz *et al*., 2021), despite its inability to chelate and mobilise iron. Scopoletin is mainly described for its selective antimicrobial activity against fungi and commensal bacteria (Ba *et al*., 2017; Harbort *et al*., 2020; Yu *et al*., 2021), but its potential role in plant iron nutrition remains undefined. Here, we show that scopoletin and the fungal root endophyte *Macrophomina phaseolina* F80 cooperate to provide mobile Fe^3+^ for subsequent import via the plant reductive import module consisting of the ferric chelate reductase FRO2 and the ferrous iron importer IRT1 (Figure 5). Our results support a pivotal role of scopoletin in the intricate interplay between F80 and its host plant, whereby scopoletin production and interaction with F80 in the rhizosphere enables the rescue of iron-deficient *Arabidopsis*.

**Fig. 5.**
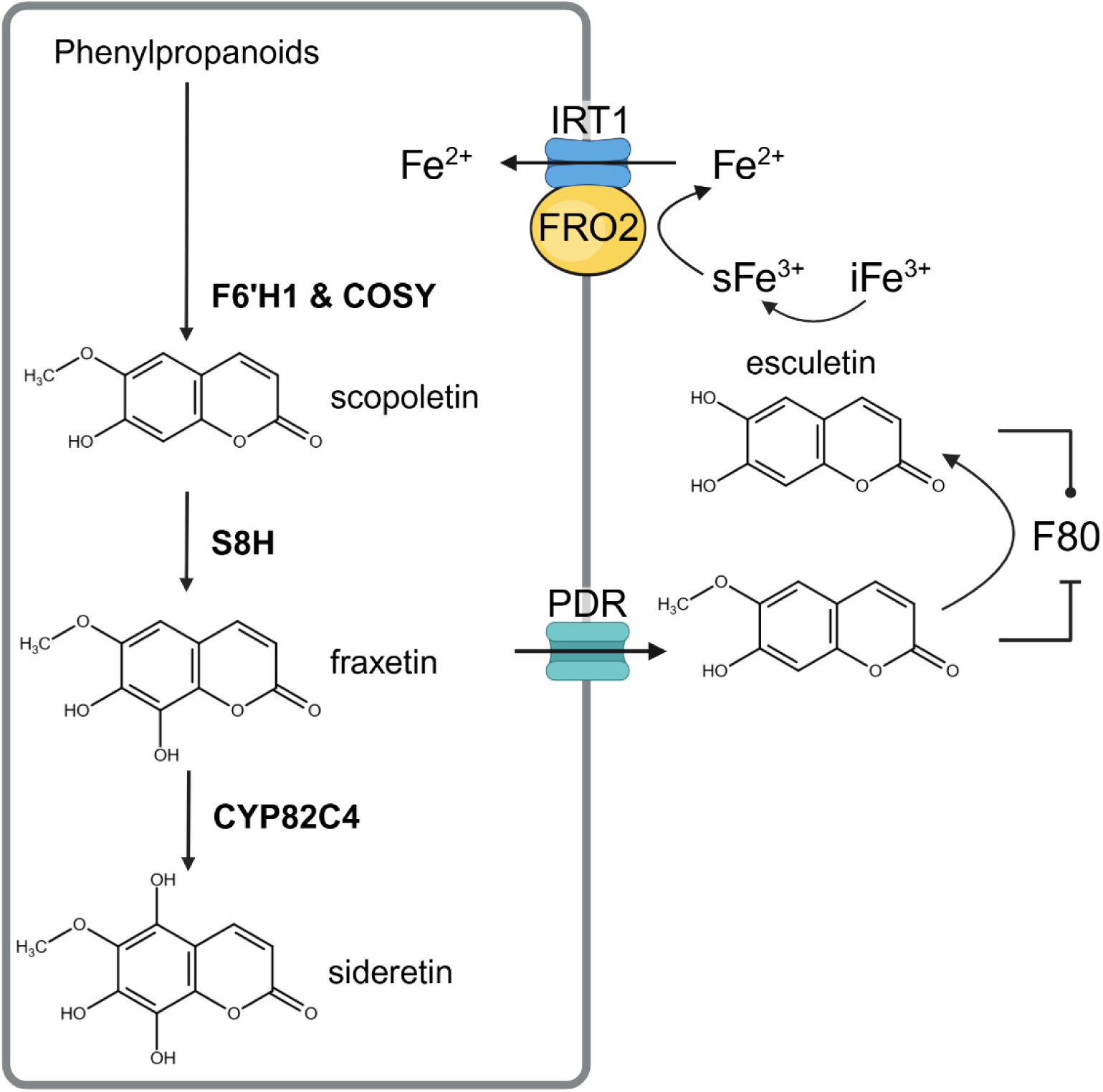
Model for the interaction of the fungal root endophyte *M. phaseolina* F80 with root-derived scopoletin to mediate iron nutrition in *Arabidopsis*. Iron stress promotes coumarin biosynthesis and exudation in *Arabidopsis* roots. The first steps in coumarin biosynthesis are catalysed by F6’H1 and COSY leading to the production of scopoletin. Subsequent conversion to fraxetin and sideretin occurs through the activity of S8H and CYP82C4, respectively. Coumarins are secreted into the rhizosphere by transporters from the PDR family. Once secreted to the rhizosphere, coumarins assist plants in iron acquisition from the soil environment. The catechol coumarins fraxetin, sideretin and esculetin can directly mobilise iron. Our model proposes a cooperative interaction between the fungal root endophyte F80 and the coumarin scopoletin, which enhances ferric iron mobilisation in the rhizosphere, upstream of the plant’s FRO2-IRT1 reductive iron uptake system. *Ex planta* data suggest that F80 mediates the conversion of scopoletin into the catechol coumarin esculetin, facilitating iron solubility and rescuing iron-deficient plant growth. (Figure made with Biorender)

Our data suggest that ferric iron mobilisation is conferred through fungal conversion of the F80-inhibitory non-catechol coumarin scopoletin to a non-inhibitory catechol coumarin esculetin which is a potent Fe^3+^ chelator (Figs 3, S3, 5) (Schmidt *et al*., 2014; Schmid *et al*., 2014; Paffrath *et al*., 2023). Because fungal mediated scopoletin conversion also occurs in the rhizosphere during plant colonisation (Fig 4c), it is likely to represent a key mechanism by which the fungus enhances iron availability. The scopoletin to esculetin transition not only enhances iron solubility but might also represent a “win–win” scenario benefiting both the fungal endophyte and its host: the fungus mitigates scopoletin antimicrobial inhibitory effects while the plant gains access to more bioavailable iron. This scenario is supported by i) *ex-planta* time-course experiments that showed decreasing abundance of scopoletin and esculetin generation in the presence of F80 and ii) cooperative iron mobilisation by scopoletin and F80 yielding similar levels of mobilised iron as those provided by esculetin alone (Fig. 3). Host rescue from limiting iron was entirely dependent on coumarin production as no rescue was observed in the coumarin-deficient *f6’h1* mutant (Fig. 2). By contrast, *s8h* mutants which produce scopoletin but lack fraxetin and sideretin supported fungal-mediated rescue of iron deficiency (Fig. 2). Thus, scopoletin appears to be necessary and sufficient for plant iron nutrition in the presence of this fungal root endophyte. A role of scopoletin in iron mobilisation was further confirmed by chemical complementation experiments. Scopoletin supplementation restored F80-mediated rescue of *f6’h1* plants to similar levels of catechol coumarins or the synthetic iron chelator EDTA in mock conditions (Fig. 4). Collectively, these findings highlight a new type of plant-microbe interaction in which a fungal endophyte repurposes a plant-derived coumarin precursor to generate bioavailable iron-chelating compounds.

Coumarin production and exudation are highly induced upon iron starvation (Fourcroy *et al*., 2014; Schmid *et al*., 2014; Sisó-Terraza *et al*., 2016; Tsai *et al*., 2018; Paffrath *et al*., 2023). Current literature suggests that fraxetin and sideretin are the primary coumarins responsible for iron acquisition (Rajniak *et al*., 2018; Tsai *et al*., 2018; Robe *et al*., 2021b; Paffrath *et al*., 2023). Based on their exudation profiles and ability to chelate and reduce iron in varying pH conditions, fraxetin appears to be more important at alkaline pH whereas sideretin primarily functions at more acidic pH (Paffrath *et al*., 2023). Scopoletin, although not an efficient iron mobiliser is, together with the redox-active sideretin the most abundant coumarin detected in *Arabidopsis* root exudates under acidic conditions (Sisó-Terraza *et al*., 2016; Rosenkranz *et al*., 2021; Paffrath *et al*., 2023). Thus, a chief role of scopoletin in shaping the rhizosphere microbiome was thought to be through its inhibitory properties (Stringlis *et al*., 2018). Our results demonstrate that scopoletin can also contribute indirectly to iron nutrition as a precursor for iron-chelating coumarins generated by microbial conversion.

F80 was isolated from apparently healthy *Arabidopsis* plants in natural populations sampled in Europe (Durán *et al*., 2018; Thiergart *et al*., 2020). Previous studies demonstrated that this fungal strain alleviated phosphate starvation in agar-plate assays (Mesny *et al*., 2021), suggesting that F80 may also be relevant in mitigating other forms of nutrient stress. These and our data reinforce the notion that F80 can exist as a neutral or beneficial endophyte associated with *Arabidopsis* roots. Conversely, *M. phaseolina* strains are well-documented plant pathogens responsible for charcoal rot, a disease affecting over 500 plant species including crops such as maize, soybean, and canola (Su *et al*., 2001; Bandara *et al*., 2018; Sinha *et al*., 2022). The ability of *M. phaseolina* F80 to degrade scopoletin might have ecological implications, for example, in promoting plant iron nutrition in certain environments and/or conferring microbial competition for niche establishment. Esculetin, for example, was found to inhibit certain fungal pathogens (Beesley *et al*., 2023). Notably, scopoletin has been implicated in plant defence as its increased accumulation was associated with enhanced resistance against pathogens (Zamioudis *et al*., 2014). Therefore, fungal-mediated degradation of scopoletin could benefit pathogens by mitigating plant defence (Sun *et al*., 2014; Gao *et al*., 2024). At this stage, it remains unclear whether scopoletin conversion by F80 is driven by a specific enzymatic activity. Alternatively, the fungus might generate a physicochemical environment that facilitates non-enzymatic oxidation and demethylation to esculetin, as previously observed during coumarin-mediated iron-oxide dissolution at the surface of iron minerals (Baune *et al*., 2020). Scopoletin was found to be degraded in four different unsterilised soils, indicating that this trait might be ecologically relevant and prevalent (Galán-Pérez *et al*., 2021).

Microbe-assisted nutrient acquisition is an emerging concept in plant nutrition, with many studies focusing on bacterial siderophores as key contributors to plant iron uptake (Kramer *et al*., 2019). In this context, the role of fungal endophytes remains largely unexplored. To our knowledge, this study provides the first mechanistic basis for an endophytic fungus directly enhancing plant iron nutrition through its interaction with a plant-derived coumarin. It therefore represents a significant advance in our understanding of how commensal fungal associations can strengthen plant nutritional resilience and productivity. Given the widespread occurrence of endophytes in natural and agricultural ecosystems and scopoletin-secreting dicot plants, future research might identify similar coumarin transformations in other plant-microbe associations.

In conclusion, we reveal here a fungal endophyte contribution to plant iron nutrition. By demonstrating that *M. phaseolina* (F80) converts scopoletin into bioactive iron-chelating coumarins, this study expands our understanding of how plant–microbe interactions influence nutrient acquisition. Fungal endophytes such as F80 could be valuable allies in enhancing plant resilience to nutrient stress, particularly in iron-deficient soils. Alongside the role of bacteria in iron nutrition (Harbort *et al*., 2020) our findings highlight the cross-kingdom significance of coumarins in microbiota-mediated plant iron acquisition. The insights gained might have implications for developing sustainable agricultural practices that leverage beneficial microbial partnerships to enhance plant nutrition and stress resilience.

## Acknowledgements

The research was funded by Deutsche Forschungsgemeinschaft SPP 2125 DECRyPT to JEP. We thank Jaqueline Bautor (Max Planck Institute for Plant Breeding Research) for experimental support, Yudelsy A. Tandron Moya (Leibniz Institute of Plant Genetics and Crop Plant Research) for conducting ICP-MS analysis, and Paul Schulze-Lefert for helpful discussions.

## Author contributions

LVD conceived the study with AP and JEP. LVD designed and performed the experiments and analysed the data. CH performed and analysed the ferrozine iron mobilisation assay. DE performed and analysed metabolite profiling. MM provided materials, analytical tools and experimental advice. RFHG supervised the ICP-MS analyses. GUB and AT supervised metabolite profiling. LVD and JEP wrote the paper with inputs from all authors.

## Data availability

Data supporting the findings of this study are available in Figs S1–S5 and Table S1.

## Supporting information

**Fig. S1** Heat-killed fungal endophyte F80 does not improve *Arabidopsis* growth under iron-limited conditions.

**Fig. S2** Phenotypes of mutants disrupted in coumarin biosynthesis or reductive iron uptake under available iron conditions in the presence or not of the fungal endophyte F80.

**Fig. S3** Changes in metabolite profile of *M. phaseolina* F80 cultures when supplemented with scopoletin.

**Fig. S4** Coumarin supplementation does not rescue *fro2* while Na_2_EDTA supplementation rescues *f6’h1*, but not *fro2*.

**Fig. S5** The concentration of mineral elements in shoots measured by ICP-MS.

**Table S1** Medium composition of ARE and vitamin solutions used in 96-Well fungal culture assay.

