## Supporting information for "Cooperation between a root fungal endophyte and host-derived coumarin scopoletin mediates *Arabidopsis* iron nutrition"

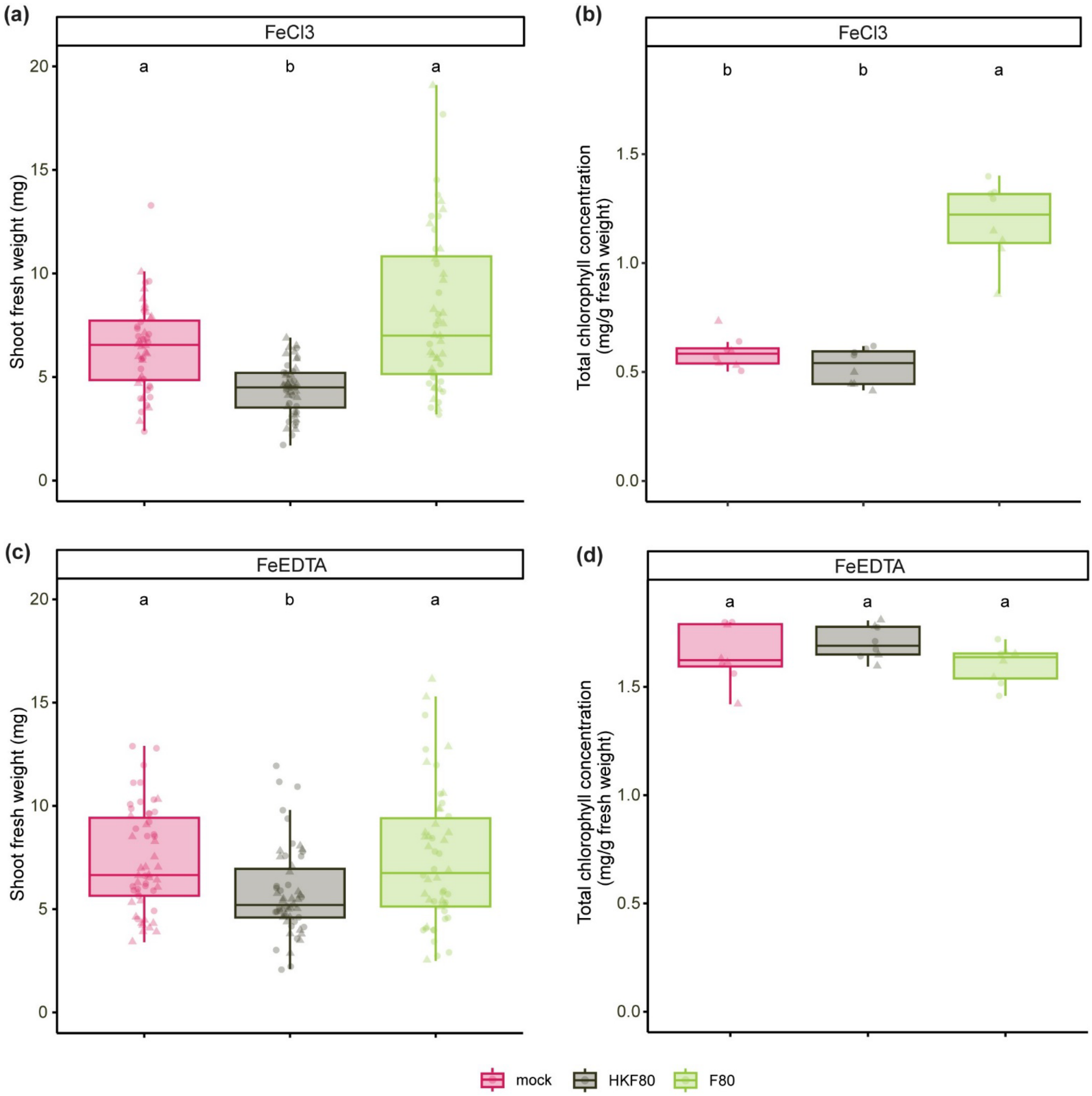


**Fig. S1 Heat-killed fungal endophyte F80 does not improve *Arabidopsis* growth under iron-limited conditions.**

(a,c) SFW and (b,d) shoot chlorophyll concentration at 2 weeks of growth after transfer. 7-day-old Col-0 seedlings were transferred to half-strength MS medium with unavailable iron (50 µM $FeCl_3$) (a,b) or available iron (50 µM FeEDTA) (c,d) at pH 5.7 mock or inoculated with heat-killed (HK) or live F80. Letters indicate significant pairwise differences between groups ($p$-adj≤0.05) by a Dunn pairwise comparison test with Benjamini-Hochberg correction. Data are from two full factorial replicates (represented by different shapes).

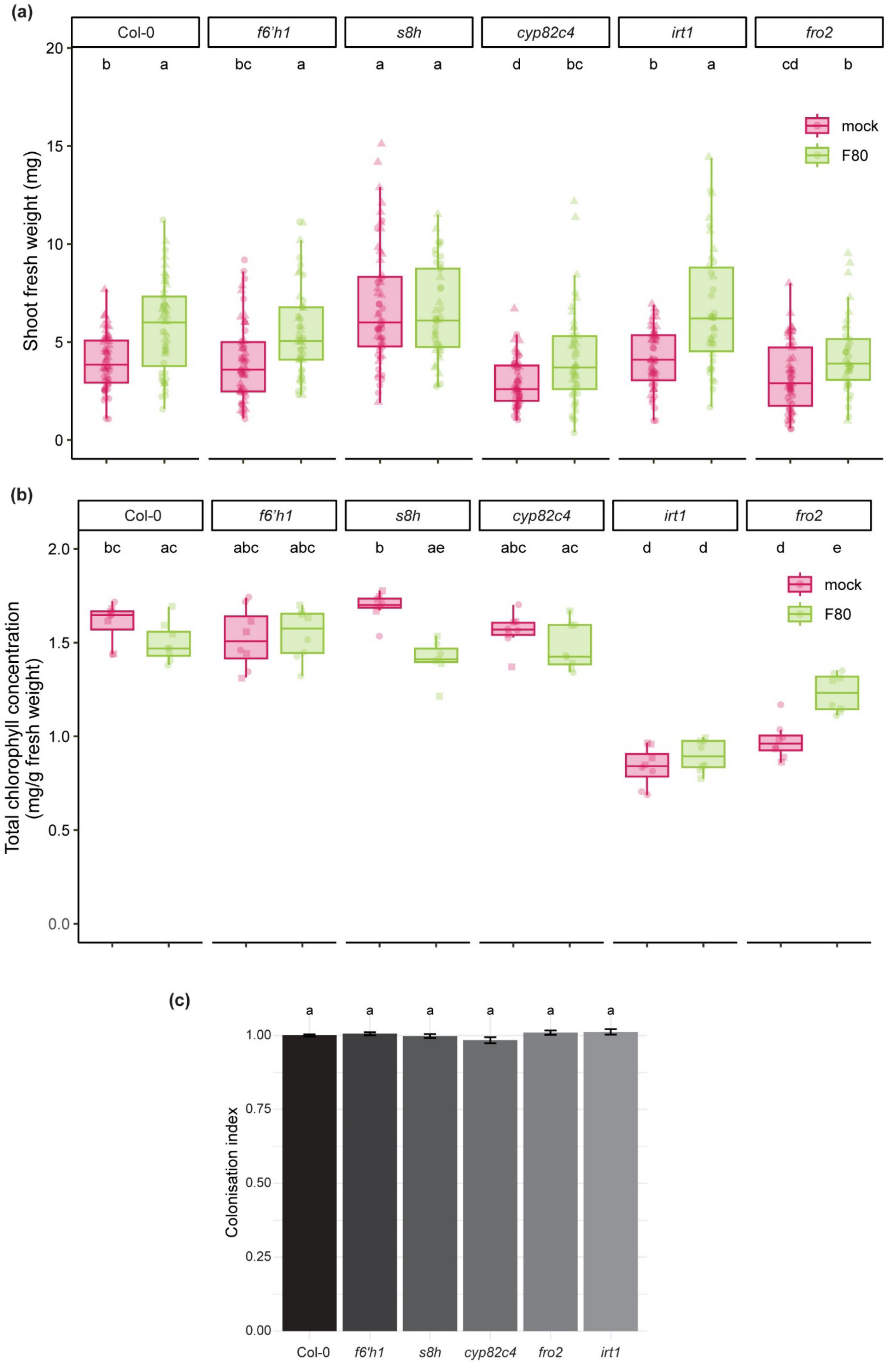
(a)
Col-0
f6'h1
s8h
cyp82c4
irt1
fro2
b a bc a a a d bc b a cd b
Shoot fresh weight (mg)
20
15
10
5
0
mock
F80
(b)
Col-0
f6'h1
s8h
cyp82c4
irt1
fro2
bc ac abc abc b ae abc ac d d d e
Total chlorophyll concentration (mg/g fresh weight)
2.0
1.5
1.0
0.5
0.0
mock
F80
(c)
a a a a a a
Colonisation index
1.00
0.75
0.50
0.25
0.00
Col-0
f6'h1
s8h
cyp82c4
fro2
irt1

**Fig. S2 Phenotypes of mutants disrupted in coumarin biosynthesis or reductive iron uptake under available iron conditions in the presence or not of the fungal endophyte F80.**

(a) SFW and (b) shoot chlorophyll content at 2 weeks of growth after transfer of indicated mutants in the coumarin biosynthesis pathway and the iron reductive import mechanism. 7-day-old seedlings were transferred to half-strength MS medium with available iron (50 µM FeEDTA) at pH 5.7 mock or inoculated with F80. Letters indicate significant pairwise differences between groups ($p$-adj≤0.05) by a Dunn pairwise comparison test with Benjamini-Hochberg correction. Data are from two full factorial replicates (represented by different shapes). (c) F80 colonisation index normalised to Col-0 control, corresponding to experiment shown in Figure 2b,c. Letters indicate significant pairwise differences between groups ($p$-adj≤0.05) by a Tukey's HSD corrected for multiple comparisons.

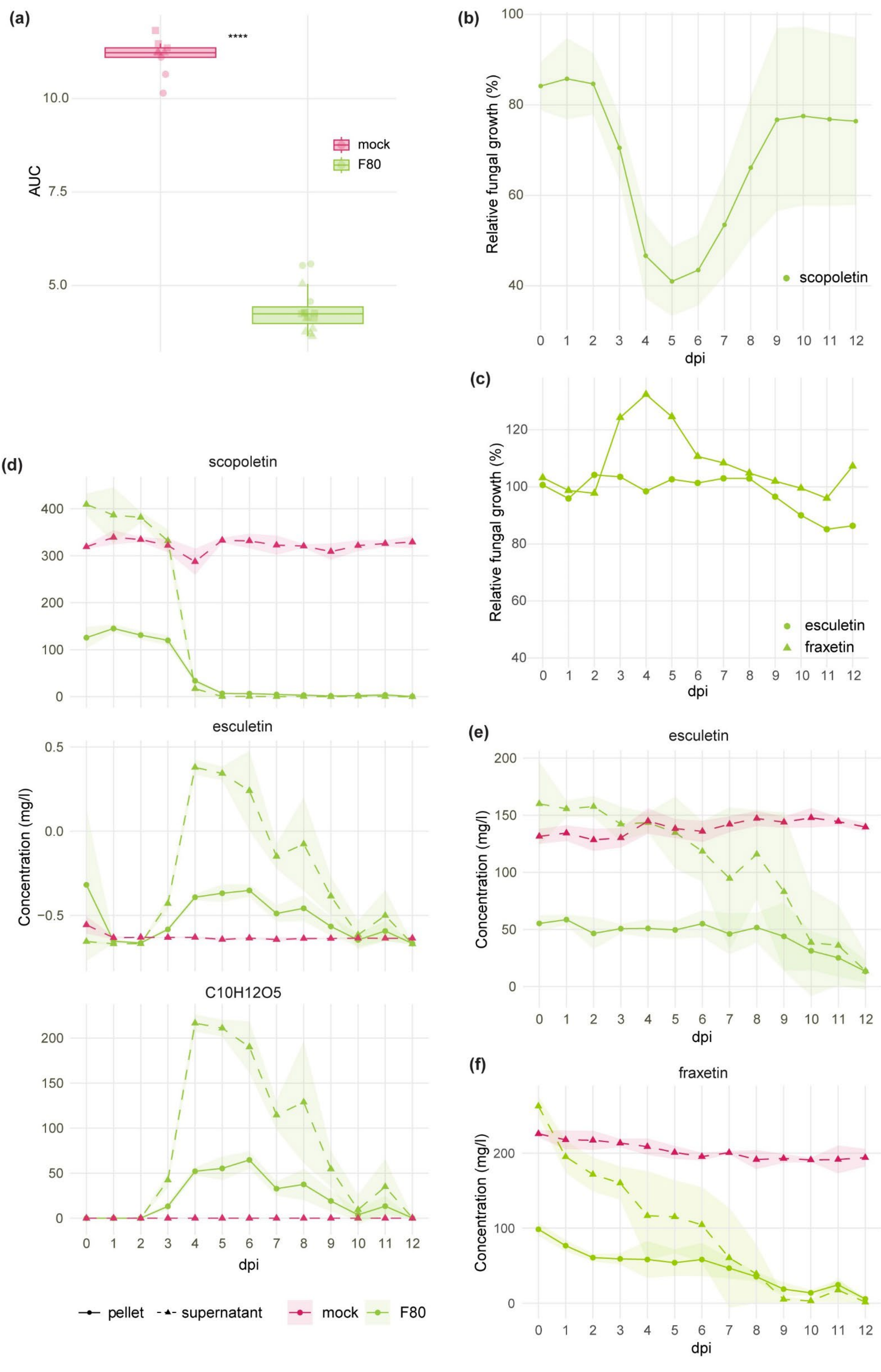
(a)
****
AUC
10.0
7.5
5.0
mock
F80
(b)
Relative fungal growth (%)
100
80
60
40
scopoletin
0 1 2 3 4 5 6 7 8 9 10 11 12
dpi
(c)
Relative fungal growth (%)
120
100
80
60
40
esculetin
fraxetin
0 1 2 3 4 5 6 7 8 9 10 11 12
dpi
(d)
scopoletin
400
300
200
100
0
esculetin
0.5
0.0
-0.5
Concentration (mg/l)
C10H12O5
200
150
100
50
0
0 1 2 3 4 5 6 7 8 9 10 11 12
dpi
pellet
supernatant
mock
F80
(e)
esculetin
Concentration (mg/l)
200
150
100
50
0
0 1 2 3 4 5 6 7 8 9 10 11 12
dpi
(f)
fraxetin
Concentration (mg/l)
200
100
0
0 1 2 3 4 5 6 7 8 9 10 11 12
dpi

**Fig. S3 Metabolite profiles of *M. phaseolina* F80 cultures when supplemented with scopoletin.** (a) Area Under the Curve (AUC) of the scopoletin detection plot in Figure 3A. Specific scopoletin fluorescence measured in *in vitro* mock and F80 cultures supplemented with 2 mM scopoletin (excitation at 385nm and emission detected at 470nm). (b) The mean OD600 of F80 cultures in 2 mM scopoletin relative to the OD600 of cultures in the absence of scopoletin from 0-12 dpi. Data combined from 3 independent biological experiments. The curve-shade indicates the standard error. (c) The mean OD600 of F80 cultures in 2 mM esculetin (circles) or fraxetin (triangles) relative to the OD600 of cultures in the absence of coumarins from 0-12 dpi. Data combined from 2 independent biological experiments. (d) Absolute quantification of the MS-TOF-IDA-MS/MS peak area data shown in Figure 3c of the indicated compounds in F80 culture supernatant (dashed line) and the fungal pellet (full line) grown for 0-12 dpi supplemented with 2 mM scopoletin. Curve-shade indicates the standard deviation. (e) Absolute quantification of the MS-TOF-IDA-MS/MS peak area data shown in Figure 3c of esculetin in F80 culture supernatant (dashed line) and the fungal pellet (full line) grown for 0-12 dpi supplemented with 2 mM esculetin. Curve-shade indicates the standard deviation. (f) Absolute quantification of the MS-TOF-IDA-MS/MS peak area data shown in Figure 3d of fraxetin in F80 culture supernatant (dashed line) and the fungal pellet (full line) grown for 0-12 dpi supplemented with 2mM fraxetin. Curve-shade indicates the standard deviation.

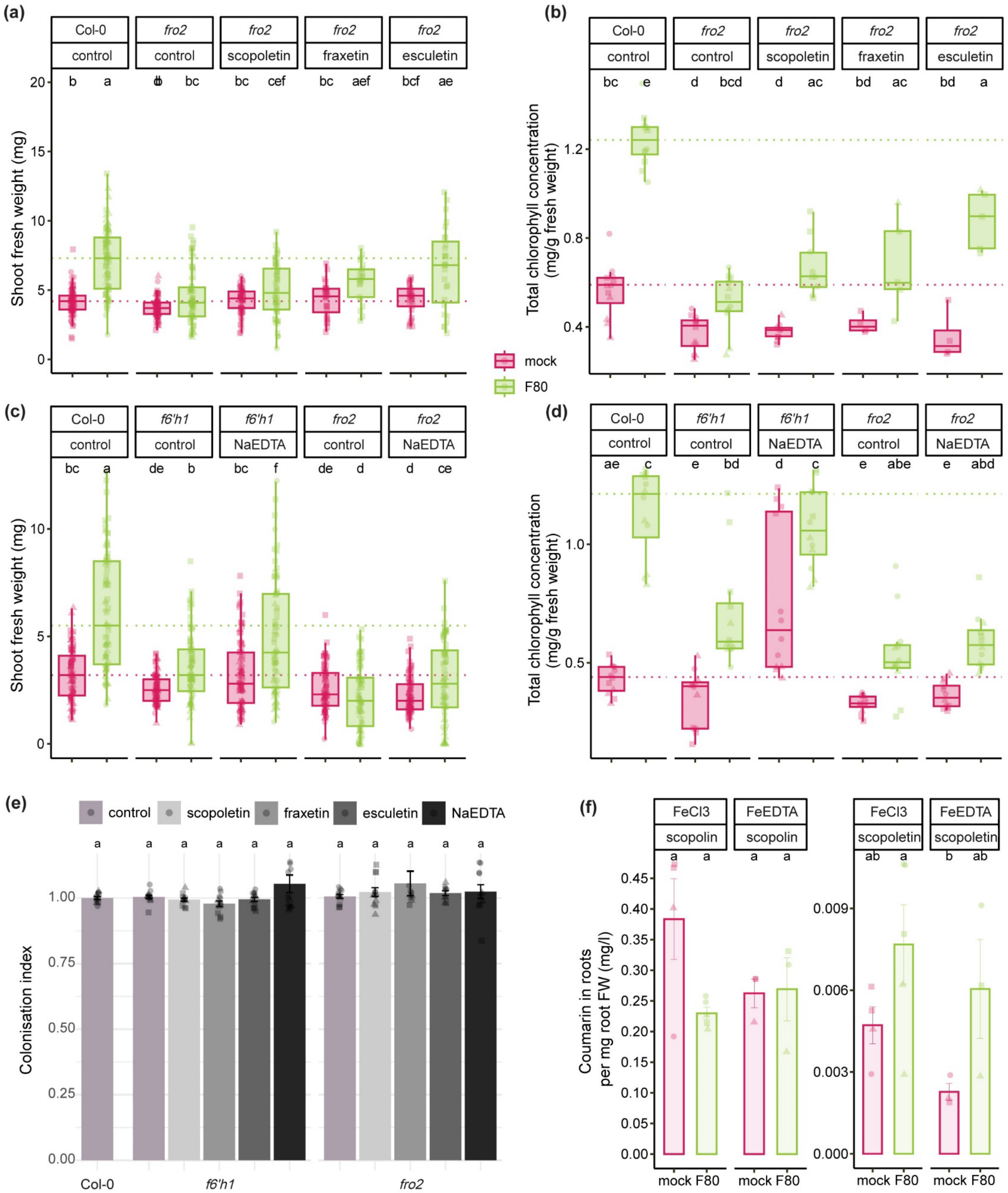


**Fig. S4 Coumarin supplementation does not rescue *fro2* while $Na_2EDTA$ supplementation rescues *f6'h1*, but not *fro2*.**

(a,c) SFW and (b,d) shoot chlorophyll concentration at 2 weeks of growth after transfer. 7-day-old Col-0, *f6'h1* and *fro2* seedlings were transferred to half-strength MS medium with unavailable iron (50 µM $FeCl_3$) at pH 5.7, supplemented with 10 µM of scopoletin, fraxetin, esculetin or $Na_2EDTA$ (indicated above the graphs) or an equal amount of DMSO (control), mock or inoculated with F80. Dashed line indicates the mean of Col-0. Letters indicate significant pairwise differences between groups (*p*-

adj≤0.05) by a Dunn pairwise comparison test with Benjamini-Hochberg correction. Data are from three full factorial replicates (represented by different shapes). (e) F80 colonisation index normalised to Col-0, corresponding to experiment shown in Figure 4 (a,b) and S4 (a,b,c,d). Letters indicate significant pairwise differences between groups (*p*-adj≤0.05) by a Tukey's HSD corrected for multiple comparisons. (f) The concentration of scopoline and scopoletin in roots 1 week past transfer of 9-day-old seedlings to half-strength MS medium with unavailable iron (50 μM FeCl3) or available (50 μM FeEDTA) at pH 5.7 mock or inoculated with F80. Approximately 150-250 mg of roots pooled from 10 plates were used to extract metabolites and analysed with MS-TOF-IDA-MS/MS. Bars represent the mean from 3 biological replicates (n=1-2) with standard error bars. Letters indicate significant pairwise differences between groups (*p*-adj≤0.05) by a Tukey's HSD corrected for multiple comparisons.

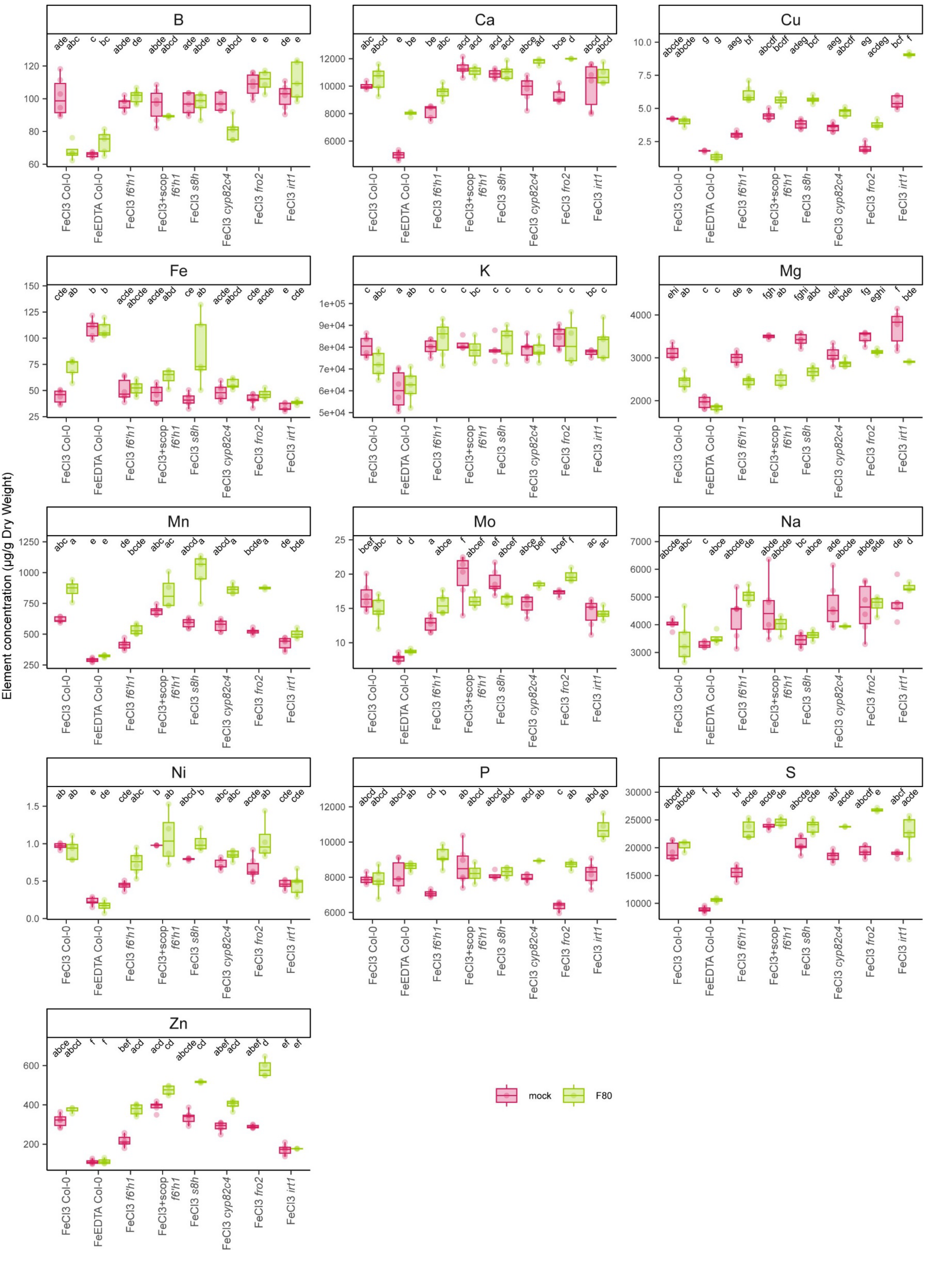

B
Ca
Cu
Fe
K
Mg
Mn
Mo
Na
Ni
P
S
Zn
Element concentration (µg/g Dry Weight)
FeCl3 Col-0
FeEDTA Col-0
FeCl3 f6'h1
FeCl3+scop f6'h1
FeCl3 s8h
FeCl3 cyp82c4
FeCl3 fro2
FeCl3 irt1
mock
F80

**Fig. S5 Concentrations of mineral elements in shoots measured by ICP-MS.**
Shoot mineral element concentrations at 2 weeks of growth after transfer. 7-day-old seedlings were transferred to half-strength MS medium at pH 5.7. Each plot depicts (from left to right) Col-0 grown on 50 µM $FeCl_3$, Col-0 grown on 50 µM FeEDTA, *f6'h1* grown on 50 µM $FeCl_3$, *f6'h1* grown on 50 µM $FeCl_3$ supplemented with 10 µM scopoletin, *s8h* grown on 50 µM $FeCl_3$, *cyp82c4* grown on 50 µM $FeCl_3$, *fro2* grown on 50 µM $FeCl_3$ and *irt1* grown on 50 µM $FeCl_3$. Letters indicate significant pairwise differences between groups ($p$-adj≤0.05) by a Tukey's HSD corrected for multiple comparisons when the data were normally distributed or by a Dunn pairwise comparison test with Benjamini-Hochberg correction when not normally distributed. Data are from three full factorial replicates.

**Table S1 medium composition of ARE and vitamin solutions used in the 96-Well fungal culture assay.**

| **5X ARE (artificial root exudates)** | |
|---|---|
| **compound** | **g/l** |
| Glucose | 8.2 |
| Fructose | 8.2 |
| Saccharose | 4.2 |
| Citric Acid | 3.2 |
| Lactic Acid | 3.2 |
| Succinic Acid | 4.6 |
| Alanine | 4 |
| Serine | 4.8 |
| Glutamic Acid | 4 |
| **100X Vitamins** | |
| **compound** | **mg/l** |
| thiamine-HCl | 10.12 |
| nicotinic acid | 12.31 |
| folic acid | 1.99 |
| pyridoxine hydrochloride | 20.56 |
| 4-aminobenzoic acid | 4.11 |
| Calcium D pantothenate | 4.77 |
| biotin | 1.00 |
